# Analyzing Vaccine Trials in Epidemics with Mild and Asymptomatic Infection

**DOI:** 10.1101/295337

**Authors:** Rebecca Kahn, Matt Hitchings, Rui Wang, Steven Bellan, Marc Lipsitch

## Abstract

Vaccine efficacy against susceptibility to infection (VE_S_), regardless of symptoms, is an important endpoint of vaccine trials for pathogens with a high proportion of asymptomatic infection, as such infections may contribute to onward transmission and outcomes such as Congenital Zika Syndrome. However, estimating VE_S_ is resource-intensive. We aim to identify methods to accurately estimate VE_s_ when limited information is available and resources are constrained. We model an individually randomized vaccine trial by generating a network of individuals and simulating an epidemic. The disease natural history follows a Susceptible, Exposed, Infectious and Symptomatic or Infectious and Asymptomatic, Recovered model. We then use seven approaches to estimate VE_S_, and we also estimate vaccine efficacy against progression to symptoms (VE_P_). A corrected relative risk and an interval censored Cox model accurately estimate VE_S_ and only require serologic testing of participants once, while a Cox model using only symptomatic infections returns biased estimates. Only acquiring serological endpoints in a 10% sample and imputing the remaining infection statuses yields unbiased VE_S_ estimates across values of R_0_ and accurate estimates of VE_P_ for higher values. Identifying resource-preserving methods for accurately estimating VE_S_ is important in designing trials for diseases with a high proportion of asymptomatic infection.

## MAIN TEXT

In 2015, the World Health Organization (WHO) identified a list of priority pathogens with potential to cause future public health emergencies of international concern (1). The Coalition for Epidemic Preparedness Innovations (CEPI) has dedicated one billion dollars to vaccine development efforts starting with three of these: Middle East respiratory syndrome (MERS) coronavirus, Lassa virus and Nipah virus (2). These three pathogens, as well as others on the WHO’s list, such as Zika virus, have high proportions of asymptomatic or mild infection (2–7). Vaccine efficacy against susceptibility to infection (VE_S_) (8), regardless of symptom level, is an important endpoint of vaccine trials for these pathogens, as infection may contribute to onward transmission, and outcomes such as Congenital Zika Syndrome, even without primary symptoms (9–14). However, VE_S_ is resource-intensive to estimate as it requires testing all trial participants, either by periodically conducting assays for infection throughout the trial, or by serologic testing at the trial’s conclusion if natural and vaccine-derived immune responses can be distinguished. Testing trial participants is also necessary for estimating a vaccine’s efficacy against progression (VEP) to symptoms, another critical outcome measurement (8). As noted in an analysis of dengue vaccine trial results, protection against symptomatic infection may in general differ from protection against infection (and in the case of dengue, VEP may be negative due to antibody dependent enhancement) (15). It is therefore important to consider estimates of both VE_S_ and VE_P_ when analyzing trial results.

We aim to identify a methodology to accurately estimate VE_S_ and VE_P_ when only a limited amount of information is available, and resources – both time and money – are constrained. Throughout, we use “asymptomatic” synonymously with “subclinical” to mean any infection episode that does not generate sufficient symptoms to prompt testing that would reveal that the participant is currently infected with the causative pathogen.

## METHODS

We model a vaccine trial by first generating a model of a main population and a network of individuals grouped into communities, the structure of which has been described previously (see Supplementary Table 1 for parameters) (16). The model is compartmental, using deterministic (differential equation) dynamics for the main population and stochastic dynamics in the communities. We simulate an epidemic in the main population with a seasonal transmission rate that generates an epidemic curve with a shape similar to the epidemic curve of the 2015 Zika outbreak in Brazil (17). The disease is introduced into communities via infectious contact with the main population, and transmission occurs when infected individuals transmit to their susceptible connections in the communities. All susceptible individuals have a daily probability of infection from each of their infectious neighbors of 1-e^-β^, where β is the force of infection, as well as a daily external hazard of infection from the main population, which varies with the prevalence in the main population. The disease natural history follows a Susceptible-Exposed-Infectious/Symptomatic (or Infectious/Asymptomatic)-Recovered (SEI_S_(I_AS_)R) model, with estimated incubation and infectious periods of a Zika-like disease (Supplementary Table 1). Vector transmission is not directly modeled, so the serial interval of the simulated disease is shorter than that of Zika virus disease. Symptomatic and asymptomatic infections are assumed to be equally infectious, and whether an individual is infected by a symptomatic or an asymptomatic individual does not affect their probability of becoming symptomatic. The baseline parameters of the model assume 20% of those infected in both the vaccine and control groups become symptomatic, based on the estimated proportion of Zika infections that are symptomatic (9). The epidemic and the vaccine trial are simulated in both a network of individuals grouped into five relatively disconnected communities as well as in a network of individuals in one large community.

7.5% of the individuals in the communities are enrolled into a 150-day long trial, and individuals are randomized to the vaccine or control groups. All individuals enrolled into the trial are assumed to be naïve to the infection, which in practice might require serologic testing of all individuals prior to enrollment. The vaccine is leaky, meaning it reduces the probability of infection upon each exposure to an infectious individual. The daily probability of infection from vaccinees’ infectious contacts is 1-e^-β(1-VE)^, where VE is the assumed direct leaky multiplicative vaccine efficacy (8,18).

VE_S_ is estimated with seven different approaches, which are described in Table 1. Trial status (i.e. vaccine or control) is the explanatory variable for all Cox proportional hazards models, and individuals who are never infected are censored at the end of the trial. Approach 1 assumes that the time of infection is known exactly (to the day) even for asymptomatic infections and therefore would be strictly applicable only where very frequent testing is performed throughout the trial. Approach 2 assumes that infection is unobserved for asymptomatic infections, so only symptomatic infections are included in the Cox regression, and those infected asymptomatically are assumed to have survived without infection to the end of follow up. Because this latter approach leads to bias in estimating VE_S_ (see Figure 1 and Results), we consider five additional approaches.

**Table 1.** Approaches for Estimating VE_S_.

| # | Approach | Symptomatic Infections | Asymptomatic Infections | Notes |
| --- | --- | --- | --- | --- |
| 1 | Cox “perfect knowledge” | Exact day of infection known | Exact day of infection known | Requires frequent viremia monitoring throughout trial |
| 2 | Cox: symptomatic only | Exact day of infection known | Treated as non-events | Assumes same proportion of vaccinated and control cases are symptomatic |
| 3 | Relative risk estimate | Ascertained prospectively and total counted at end of trial | Serologic testing at end of trial | $\widehat{VE}_S = 1 - \frac{\text{Attack Rate (Vaccinated)}}{\text{Attack Rate (Control)}}$ |
| 4 | Corrected relative risk estimate (23) | Ascertained prospectively and total counted at end of trial | Serologic testing at end of trial | $\widehat{VE}_S = 1 - \frac{\ln(1 - \text{Attack Rate (Vaccinated)})}{\ln(1 - \text{Attack Rate (Control)})}$ |
| 5 | Interval censored Cox model (3 intervals) | Exact day of infection known | Interval for infection time known: 3 serologic tests | Interval based on serologic testing 2 times throughout trial and once at end (i.e. day 50, day 100, day 150) |
| 6 | Interval censored Cox model (1 interval) | Exact day of infection known | Interval for infection time: length of trial | Serologic testing once at end of trial |
| 7 | Imputation | Exact day of infection is known | Serologic testing at end of trial for asymptomatic participants in a random sample of 10% of participants and the remaining asymptomatic participants’ event statuses are imputed | a) Probability of infection (in 1 community analysis) or ratio of asymptomatic to symptomatic infections (in 5 community analysis) estimated in sample of 10% of the vaccinated and the control groups<br>b) Infectious status of the remaining asymptomatic individuals imputed using multiple (10) imputation (26) |
|  |  |  |  | c) Imputed data set then analyzed with Approach 6 |

**Figure 1.**
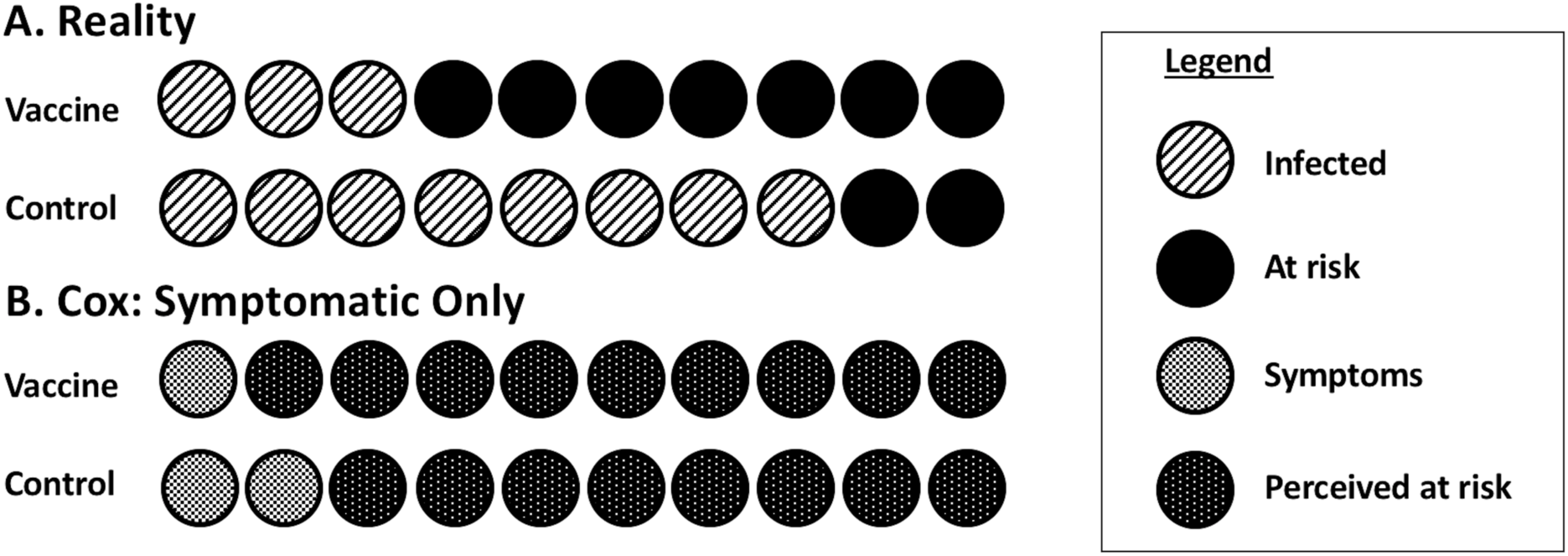
Differential misclassification of at risk person-time. When considering only symptomatic events, presumed person-time at risk increases for both the vaccine and control groups, as all asymptomatic infections are now perceived to be uninfected and at risk for the entire period of the trial. In the vaccine group, 9 people are perceived to still be at risk (Panel B), when in reality only 7 remain at risk (Panel A), as 2 people are asymptomatically infected. In the control group, 8 people are perceived to be at risk (Panel B), when in reality only 2 remain at risk (Panel A). Because there are more people infected and therefore more people incorrectly still perceived at risk in the control group than the vaccine group, apparent incidence is underestimated in the controls more so than in the vaccine group, leading to the bias towards the null. This bias is exacerbated as R_0_ increases and more people in the control group become infected but are still perceived to be at risk. At time t post randomization, person time at-risk in the controls will be overestimated by a factor 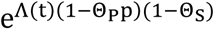 relative to the vaccine group, where Λ(t) is the cumulative hazard to time t, p is the symptomatic proportion in controls, and 1-θ_S_ and 1-θ_Ρ_ are the efficacy of the vaccine against infection, and disease given infection, respectively (27). This will be greater than 1 for non-negative VE_P_ and positive VE_S_.

In the interval censored Cox models, the exact day of infection for the symptomatic individuals is known (and in practice would be laboratory-confirmed). For the asymptomatically infected individuals, the interval for day of infection ranges from the day of their last negative serologic test to the day of their first positive serologic test. Two different interval lengths are assessed to determine if increased frequency of testing yields more precise results (19). As mentioned above, this method assumes the ability of the serologic test to distinguish between vaccine acquired immunity and naturally acquired immunity, currently possible for some but not all vaccines/pathogens (20–22).

The results from the network with five communities are analyzed first with the same seven approaches, treating the five communities as if they were one large community. Alternatively, to account for the potential for heterogeneity in hazard rates between communities, the Cox models in Approaches 1, 2, 5, 6 and 7 are stratified by community (16), and estimates from Approaches 3 and 4 are calculated separately within each community and meta-analyzed using inverse-variance weighting.

Empirical coverage probabilities are calculated by the proportion of simulations with 95% confidence intervals that cover the true VE_S_ parameter of the model (60%). Power is estimated by the proportion of simulations in which P < 0.05 for the null hypothesis of VE_S_=0 and the estimated VE_S_ is greater than 0. The trial is also simulated with fewer participants to assess power in smaller trials.

Additionally, to evaluate the efficacy of the vaccine in preventing progression to symptoms, VE_P_ is estimated by:

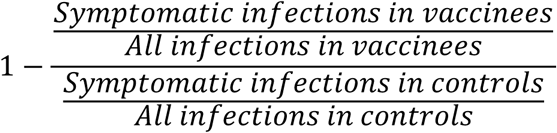

Finally, to assess whether the results hold in other contexts, trial parameters such as trial length, VE_S_, and the proportion of the infected individuals in each arm of the trial that becomes symptomatic, are varied.

R code for these analyses is available at: https://github.com/rek160/Asymptomatic-Infection-Vaccine-Efficacy.

## RESULTS

Figure 2 displays the results of the median of 500 simulations in the single-community network, showing VE_S_ estimates from the seven approaches described above across three values of R_0_, the basic reproductive number. As expected, Approach 1 returns accurate VE_S_ estimates, while Approach 2 returns estimates biased towards the null because there is differential overestimation of person-time at risk, with worse overestimates in the controls (Figure 1). This bias is exacerbated as R_0_ increases. Approach 3 returns an estimate biased toward the null compared to the true value of VE_S_, also as expected and also worsened at higher levels of R_0_ (8). Approach 4 corrects this bias by converting the risk-based analysis into a rate-based analysis (18,23).

**Figure 2.**
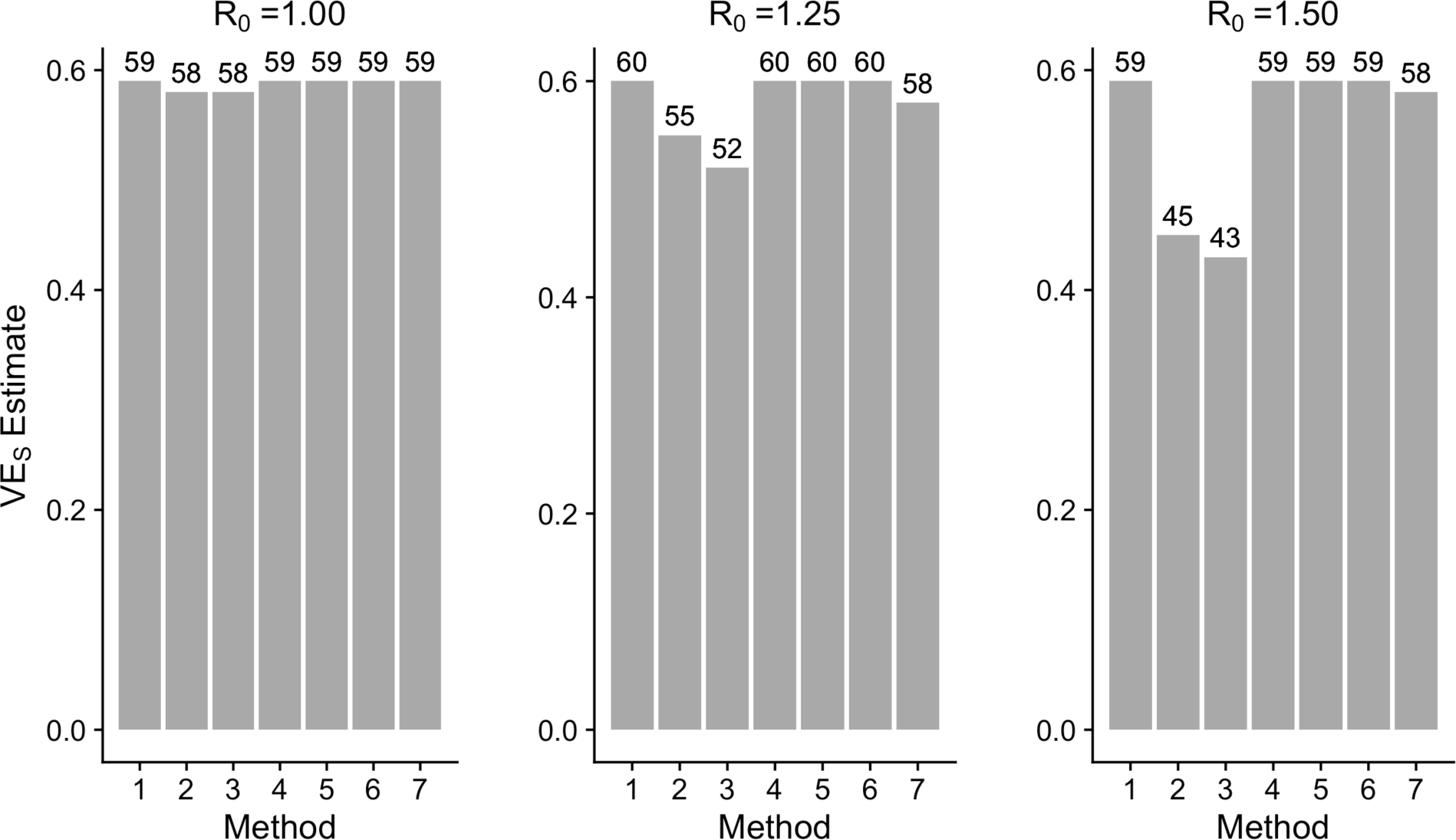
VE_S_ estimates (baseline parameters, one community network). Estimates for vaccine efficacy against susceptibility to infection (VE_S_) using seven different approaches across three values of R_0_ under the model’s baseline parameters in the one community network.

The interval censored Cox proportional hazards models (Approaches 5-6) return estimates approximately equal to the VE_S_ input into the model. These approaches require fewer resources than would be necessitated by the Cox model with perfect knowledge of infection time (Approach 1) because they use only three or one serologic tests, respectively, rather than frequent monitoring for infection throughout the period of follow-up. Approach 6, the interval censored Cox model with testing only at the end of the trial, yields the same results as Approach 5, testing three times, without substantial difference in coverage probability or power in the settings considered (Table 2 and Figure 3). Thus, both Approaches 4 and 6 yield accurate estimates with testing only required once at the end of the trial.

**Table 2.** VE_S_ Estimates (Empirical Coverage Probabilities)^a^.

| Method | $R_0 = 1.00$ | | $R_0 = 1.25$ | | $R_0 = 1.50$ | |
| --- | --- | --- | --- | --- | --- | --- |
|  | 1 community | 5 communities | 1 community | 5 communities | 1 community | 5 communities |
| 1 | 0.59 (0.96) | 0.59 (0.95) | 0.60 (0.95) | 0.59 (0.94) | 0.59 (0.93) | 0.59 (0.94) |
| 2 | 0.58 (0.96) | 0.58 (0.94) | 0.55 (0.90) | 0.52 (0.85) | 0.45 (0.49) | 0.46 (0.52) |
| 3 | 0.58 (0.95) | 0.58 (0.95) | 0.52 (0.51) | 0.51 (0.49) | 0.43 (0) | 0.44 (0) |
| 4 | 0.59 (0.96) | 0.59 (0.94) | 0.60 (0.95) | 0.59 (0.95) | 0.59 (0.94) | 0.59 (0.94) |
| 5 | 0.59 (0.96) | 0.59 (0.95) | 0.60 (0.94) | 0.59 (0.94) | 0.59 (0.94) | 0.59 (0.93) |
| 6 | 0.59 (0.95) | 0.59 (0.95) | 0.60 (0.94) | 0.59 (0.93) | 0.59 (0.93) | 0.59 (0.92) |
| 7 | 0.59 (0.91) | 0.59 (0.96) | 0.58 (0.92) | 0.58 (0.97) | 0.58 (0.91) | 0.58 (0.96) |

<sup>a</sup>Empirical coverage probabilities are calculated by the proportion of simulations with 95%
confidence intervals that cover the true $VE_S$ parameter of the model (60%).

**Figure 3.**
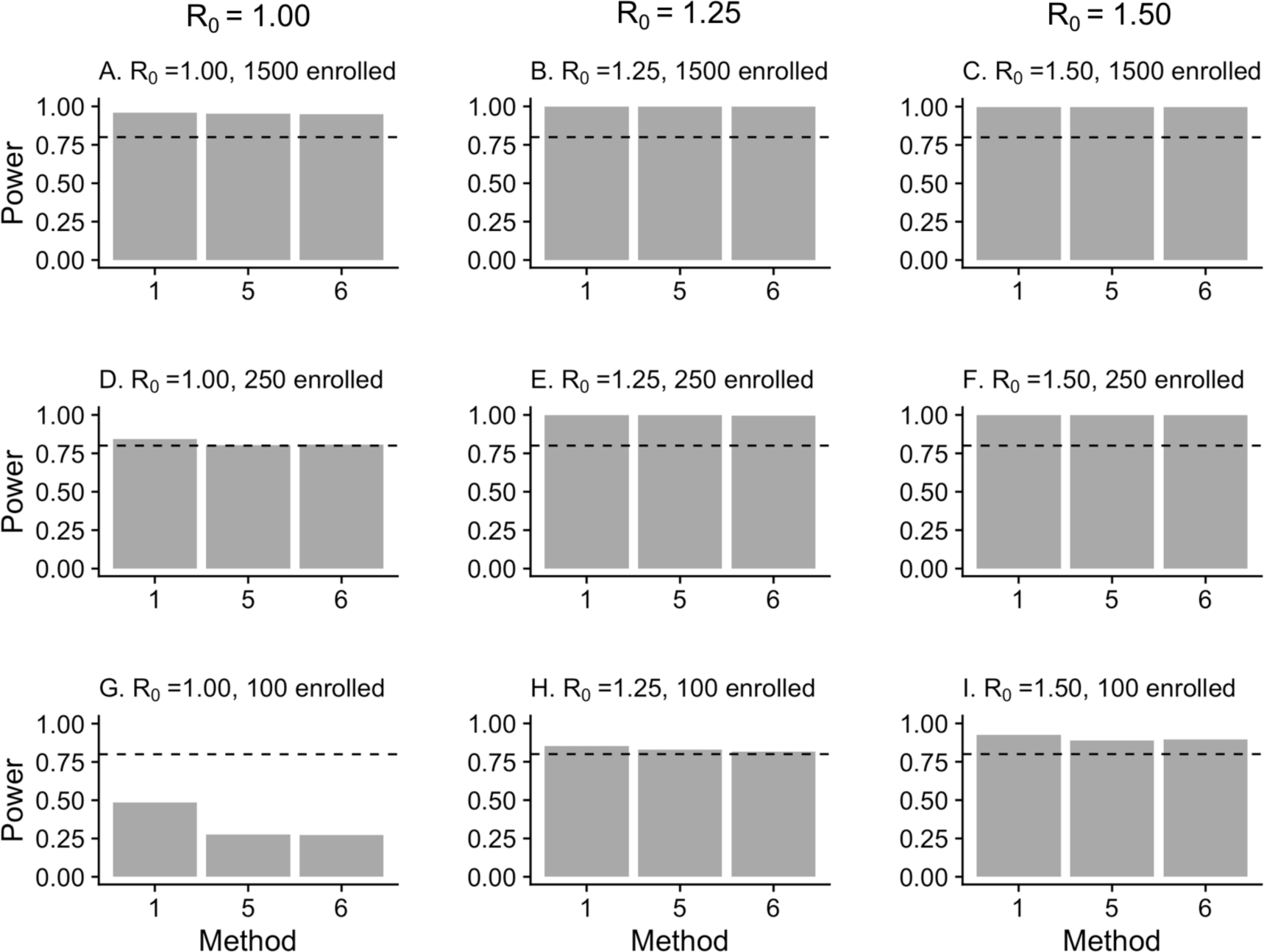
Power. The power of the Cox “perfect knowledge” and the two interval censored models in one community with A-C) 1500 trial participants (baseline), D-F) 250 trial participants, and G-I)100 trial participants across the three values of R_0_. The interval censored models do not lead to a substantial loss in power except in the trial with 100 participants enrolled when R_0_ = 1. The dashed lines are at 0.80.

Even a single serologic test could be resource-intensive. Approach 7, which only requires testing 10% of the trial participants at the end of the trial, results in accurate estimates of VE_S_ (Figure 2) for all values of R_0_ considered and of VE_P_ (Table 3) for R_0_ values of 1.25 or 1.50. Only testing 10% of the trial participants once at the end of the trial substantially reduces required resources. However, when the number of cases is very low (Supplementary Table 2), the sample does not accurately estimate VE_P_, and this approach is less efficient than others in many of the settings considered (i.e. wider confidence intervals, see Supplementary Table 3). Similar results are obtained in the analyses of the five communities (Figure. 4-5). However, when the number of cases is low (R_0_=1), the meta-analyses of Approaches 4 and 7 are imprecise.

**Table 3.** Median VE_P_ Estimate (True VE_P_ = 0) in Full Trial and Sample From Approach 7.

| $VE_P$ | $R_0 = 1.00$ | | $R_0 = 1.25$ | | $R_0 = 1.50$ | |
| --- | --- | --- | --- | --- | --- | --- |
|  | 1 community | 5 communities | 1 community | 5 communities | 1 community | 5 communities |
| Full Trial | -0.02 | 0.003 | 0.02 | -0.01 | -0.01 | -0.002 |
| Sample | 0.38 | 0.13 | 0.04 | -0.01 | -0.03 | -0.003 |

**Figure 4.**
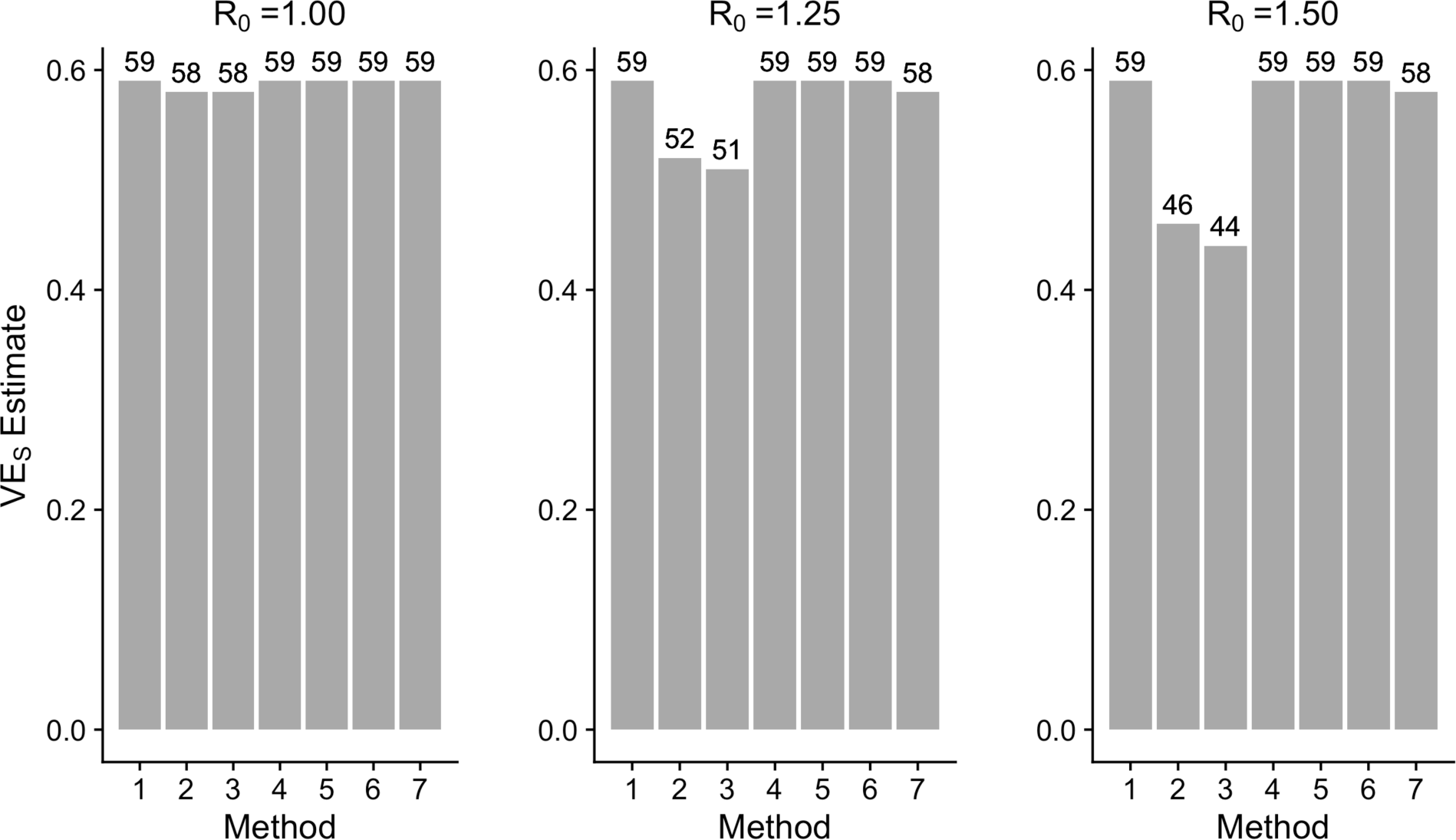
VE_S_ estimates (baseline parameters, five communities network, analyzed as one large community). Estimates for vaccine efficacy against susceptibility to infection (VE_S_) using seven different methods across three values of R_0_ under the baseline parameters in the five communities network analyzed as one large community.

**Figure 5.**
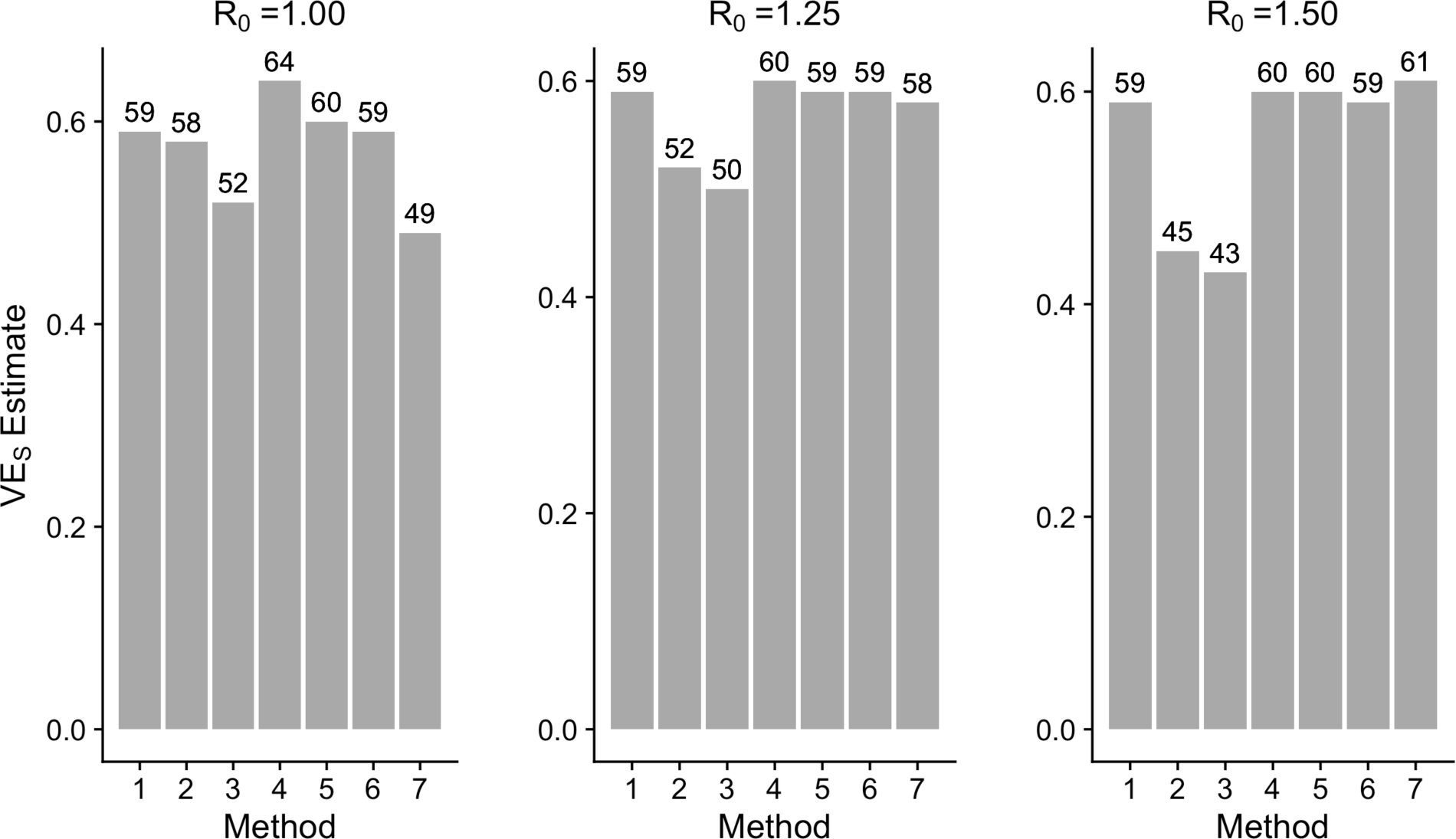
VE_S_ estimates (baseline parameters, 5 communities network, stratified analyses). The estimates for vaccine efficacy against susceptibility to infection (VE_S_) using seven different methods across three values of R_0_ under the baseline parameters in the five communities network, with stratified and meta-analyzed analyses.

Results are essentially unchanged across changes in duration of the trial or vaccine efficacy, and when the proportion symptomatic among the vaccine and control groups differs (i.e. VE_P_ ≠ 0), as shown in Figures S1-5. When the vaccine has an effect on both susceptibility to infection and progression to symptoms, serologic testing helps differentiate between VE_S_ and VE_P_ except when the number of total cases is low.

## DISCUSSION

For pathogens with a high proportion of mild or asymptomatic infection, understanding if the vaccine prevents all infection, not solely symptomatic infection, as well as understanding the vaccine’s impact on progression to symptoms, is critical for determining the epidemiologic impact of the vaccine. However, costs and resources can pose major barriers to estimating these critical values. Here we have discussed different methods and their varying levels of accuracy and resource requirements for estimating VE_S_ and VE_P_. The corrected relative risk estimate (Approach 4), the interval censored Cox models (Approaches 5-6), and the imputed interval censored Cox model (Approach 7) provide estimates close to the VE_S_ input into the model across values of R_0_, which is of course also obtained under the assumption of perfect knowledge of the time of all asymptomatic infections (Approach 1). A Cox model considering only symptomatic infections proves biased, especially at higher values of R_0_. Approaches 1 and 4-7 return accurate estimates of VE_P_, with the exception of Approach 7 when R_0_ is low due to the small number of cases overall and in the sample.

In practice, using a Cox proportional hazards model for the time of all infections would entail testing everyone frequently (perhaps weekly or even daily) for infection throughout the trial, requiring significant expenditures of both money and time. Using a corrected relative risk estimate or an interval censored Cox model, an accurate estimate of VE_S_ and VE_P_ can be obtained with serologic testing only once at the end of the trial. Testing only 10% of the trial population and imputing the event status of the remaining asymptomatic trial participants substantially reduces the resources needed while still providing critical information about the vaccine.

These approaches work in both trials conducted in one large community and trials conducted in disconnected communities, such as the recent malaria trials (24). In trials with more communities or increased heterogeneity, the bias from heterogeneity in hazard rates will likely be more pronounced (16). Methods to account for this heterogeneity, such as stratification, meta-analyses, or incorporation of random effects, should therefore be used; although when the number of cases is very low, some of these methods may be imprecise.

### Limitations

While Approaches 4-7 require substantially fewer resources for estimating VE_S_ and VE_P_ than Approach 1 and are more accurate than Approaches 2-3, all participants must be tested at the beginning of the trial to ensure they are not exposed or immune. Including those with pre-existing immunity in the trial would reduce the total number of overall cases, and thus the power, limiting the ability to draw a statistically significant conclusion about the vaccine’s effects. This challenge however is not limited to diseases with high proportions of mild or asymptomatic infection, as prior immunity would reduce the power of any trial. Additionally, distinguishing between natural and vaccine-derived immunity can be challenging, especially at the beginning of an outbreak of an emerging infectious disease about which not much is known and for which serologic tests are likely in early stages of development (22,25). Therefore, investments in diagnostic development will also be critical for ensuring accurate VE estimates. Finally, many simplifying assumptions are made, including complete ascertainment of infectious cases, perfect sensitivity and specificity of the diagnostic test, and comparability of the infected vaccinees and infected controls for the estimation of VE_P_ (8).

We have identified methods that accurately estimate VE_S_ and VE_P_ and only require serologic testing of trial participants once at the end of the trial. Only acquiring serological endpoints in a 10% sample yields unbiased VE_S_ estimates, significantly reducing required resources. While parameterized for a Zika-like disease, the methods and exact parameters described above are not meant to represent a particular epidemic context, but merely to serve as a guide when thinking through how to accurately estimate important endpoints of vaccine trials in different settings with limited resources and information. R code for this simple model can be readily modified to reflect disease-, vaccine- and setting-specific parameters. Identifying resource-preserving methods is important in designing trials for diseases with a high proportion of asymptomatic or mild infection, especially when those cases are still infectious. Understanding the potential sources of bias from different approaches can allow for more accurate estimates in epidemic settings.

